# Feline calicivirus encoding Nano Luc luciferase as a tool for assessing antibody neutralisation and antivirals

**DOI:** 10.64898/2026.08.20.745972

**Authors:** Hagar Sasvari, Kelsey Urquhart, Rahaf Alharbi, Marie McCallum, Lotta H Truyen, Sora Ogawa, Juan Barcena, Matteo Bordicchia, Vanessa R Barrs, David Bhella, William Weir, Brian J Willett, Margaret J Hosie, Lee Sherry

## Abstract

Feline calicivirus (FCV) is among the most common viruses to infect cats worldwide, with prevalence estimated to range from 10-90% depending on the population sampled. Typical FCV infection presents with oral ulcerations, fever and in some cases can also lead to clinical signs such as pneumonia or ‘limping syndrome’. However, some FCV strains have been isolated from cats exhibiting virulent systemic (VS) disease, which is associated with high morbidity and mortality. Breakthrough VS-FCV infections have been recorded in vaccinated cats and, therefore, there is considerable interest in developing novel therapeutics for use in the face of VS-FCV outbreaks. However, to design effective therapeutics, a tractable system to systematically assess the efficacy of novel vaccine candidates or antivirals is required.

Here, we used reverse genetics to develop an FCV reporter virus, inserting NanoLuc luciferase into the LC protein of FCV-Urbana (FCV-Urbana^NL^). We characterised the replication kinetics of FCV-Urbana^NL^ in comparison to its parent virus and assessed the stability of the reporter over multiple passages. Subsequently, we developed virus neutralisation assays to assess a range of monoclonal antibodies that recognise FCV Urbana. We then assessed the breadth of neutralisation by exchanging the major capsid protein, VP1, of FCV Urbana with VP1 from the vaccine strain F9 and the VS-FCV strain NSW-E1. Finally, we evaluated the utility of the FCV^NL^ reporter system to screen candidate antiviral compounds, identifying GS-441524 (the active metabolite of the parent nucleoside remdesivir) as having therapeutic potential against FCV. These findings highlight the potential of this reporter virus as a powerful molecular tool to accelerate the discovery and development of novel therapeutics.

## Introduction

Feline calicivirus (FCV) is a common virus that infects cats, with a prevalence ranging between 2.5% and 32% worldwide (1–3) that is broadly proportional to the number of cats per household; FCV prevalence can reach 90% among shelter and stray cat colonies (4). While FCV infection is sometimes inapparent, it commonly causes mild disease, presenting with a range of clinical signs that include oral ulceration, sneezing, nasal discharge and fever (5). In some cases, infection can also lead alternatively to pneumonia or ‘limping syndrome’ (6,7).

However, more severe strains of FCV causing virulent systemic (VS) disease have emerged since the identification of the first case in 1998 (8). VS-FCV is characterised by a widespread infection resulting in oedema, ulcers, fever, anorexia and multiorgan failure; 30-70% of VS-FCV cases lead to death (8,9). VS disease is of significant concern for the cat population as it cannot be differentiated from other infections early on and there is a lack of effective treatment and preventive options (8,10). Although there are effective FCV vaccines available, immunisation does not protect cats from VS-FCV infection or severe disease (4).

FCV belongs to the *Vesivirus* genus of the *Caliciviridae* family, which is characterised by a single-stranded positive sense RNA genome (11). The FCV genome is approximately 7.7 kb in size and comprises 3 distinct open reading frames (ORFs). ORF 1 is translated from the genomic RNA and encodes the non-structural proteins (NS1-NS7), which are translated as a large polyprotein precursor that is co-and post-translationally processed into mature viral proteins by the viral protease/polymerase complex, NS6/7. During infection, FCV produces a subgenomic RNA, which encodes the viral capsid proteins VP1 and VP2. ORF2 is initially translated as a precursor protein comprised of the leader of the capsid protein (LC) and VP1, which is post-translationally cleaved producing the major capsid protein, VP1, and LC; the latter has been hypothesised to aid in viral release through the induction of apoptosis (4). ORF3 encodes the minor capsid protein VP2, which, despite its lower abundance relative to VP1, is critical for the production of infectious virus particles (12–14).

Infectious FCV virions are ∼38 nm diameter icosahedral capsids that consist of 180 copies of VP1 and only 12 copies of VP2 (4,13). To facilitate viral entry, the FCV capsid binds to the feline junctional adhesion molecule 1 (fJAM-A) functional receptor and then enters the cell through clathrin-mediated endocytosis. The interaction of the viral capsid with fJAM-A triggers conformational changes within the virion leading to the formation of a large-portal like structure formed by the 12 VP2 proteins present within the virion (13). The VP2 portal penetrates the endosomal membrane to release the genome, with the highly conserved hydrophobic amino acids at the N-terminus of the VP2 protein playing a key role in this process (14).

Traditionally, neutralising antibody activity is measured using plaque reduction neutralisation tests (PRNTs), which remain the reference standard for serological testing and determining immune correlates of protection (15). Similarly, in screening tests for small molecule inhibitors of viral infection, compounds are often added during infection and end-point viral titres are assessed by plaque assay to determine compound efficacy (16). However, such low throughput assays are not optimal for large-scale serological studies, nor for antiviral development. Therefore, reporter viruses have been developed across a number of different virus families, increasing throughput and assay sensitivity whilst yielding results equivalent to reference PRNTs (16–19).

In this study, we employ reverse genetics to build on previously reported fluorescent FCV reporter viruses (20) to develop and characterise a FCV NanoLuc luciferase (NanoLuc) reporter virus system that provides rapid readouts for viral replication kinetics and increased sensitivity for the assessment and development of monoclonal antibody (mAb) and antiviral compounds.

## Results

### Development and characterisation of a NanoLuc-encoding FCV

NanoLuc reporter viruses are highly effective tools for assessing *in vitro* and *in vivo* viral replication, antiviral immunity and the impact of novel antivirals (21–23). However, no similar system has been described for FCV. Here, we build on previous work (20) by replacing the fluorescent protein reporter with NanoLuc to produce a highly sensitive reporter virus for FCV. To generate a replication-competent NanoLuc reporter virus, the NanoLuc sequence was inserted into the Leader of the Capsid (LC) region of FCV-Urbana^wt^ to generate FCV-Urbana^NL^ (Figure 1A). To assess the impact of inserting NanoLuc into FCV-Urbana on viral replication kinetics, CRFK cells were infected at a multiplicity of infection (MOI) 1 with either FCV-Urbana^wt^ or FCV-Urbana^NL^ and supernatants were collected at 6, 12, 24, 32 and 48 hours post-infection (hpi). FCV-Urbana^NL^ displayed diminished viral replication kinetics in comparison to FCV-Urbana^wt^ with a peak viral titre reduction of ∼1.5 log_10_. However, FCV-Urbana^NL^ showed exponential growth, similar to that of the parental virus (Figure 1B).

**Figure 1:**
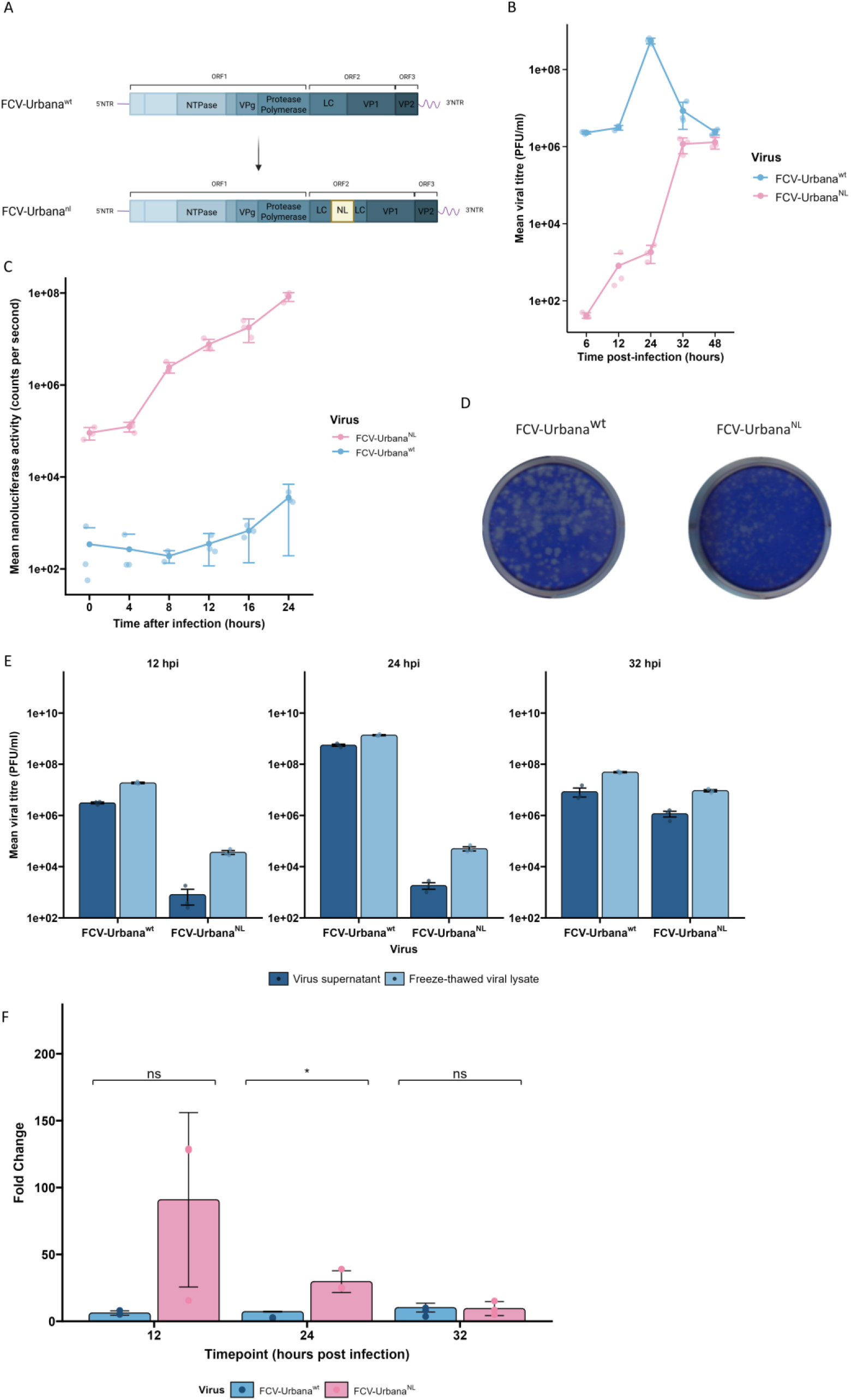
Recovery of FCV NanoLuc Urbana (FCV-Urbana^NL^) alongside wild-type FCV-Urbana (FCV-Urbana^wt^). (A) Schematic of FCV virion structure, showing the placement of capsid proteins VP1 and VP2. In FCV^NL^ viruses, NanoLuc is inserted into the Leader of the Capsid region of ORF2. (B) Comparison of viral replication kinetics between FCV-Urbana^wt^ and FCV-Urbana^NL^ following infection at MOI 0.1 (C) NanoLuc activity elicited by infection with FCV-Urbana^wt/NL^ at 100 PFU/mL. (D) Plaque assay showing differences in viral plaque morphology for FCV-Urbana^wt^ and FCV-Urbana^NL^, alongside corresponding titres of newly rescued virus stocks. (E and F) Virus stocks were subjected to three freeze-thaw cycles, alongside non-freeze-thawed controls to investigate potential for attenuation after LC-insertion; statistical significance was determined by student’s t test with results denoted with * P < 0.5, and ns for not significant differences. (n=3)

To assess the functionality and specificity of the FCV-Urbana^NL^ virus, CRFK cells were infected at 100 plaque forming units per well (PFU/well) with either FCV-Urbana^wt^ or FCV-Urbana^NL^ and NanoLuc activity was measured at different timepoints (0, 4, 8, 12, 16, 24 hpi). FCV-Urbana^NL^ produced significantly higher luminescence outputs, with luciferase activity first detected between 4 and 8 hpi, displaying a 3-log_10_ difference compared to FCV-Urbana^wt^ at 8 hpi, increasing to 4-log_10_ difference by 24 hpi (Figure 1C). Therefore, these data highlight that the presence of NanoLuc provides a sensitive and specific readout for viral replication in CRFK cells.

To further evaluate the impact of the NanoLuc insertion on viral replication kinetics, we compared the plaque morphology of FCV-Urbana^wt^ and FCV-Urbana^NL^ (Figure 1D). FCV-Urbana^wt^ formed large plaques, while FCV-Urbana^NL^ plaques were notably smaller, consistent with attenuation of the virus. This attenuation likely results from the NanoLuc sequence being inserted into the LC region, which has been shown previously to be responsible for viral egress. To evaluate the impact of inserting NanoLuc into LC on viral egress, CRFK cells were infected with either FCV-Urbana^wt^ or FCV-Urbana^NL^ at MOI 1 and 500 µL viral supernatants were collected at each time-point, and then the remaining supernatant and cell monolayer was subjected to three freeze-thaw cycles (Figure 1E). The resulting samples were subsequently titrated by plaque assay. While FCV-Urbana^wt^ displayed modest increases in viral titre following freeze-thaw throughout the time-course, FCV-Urbana^NL^ showed increases over 1 log_10_ following freeze-thaw cycles at 12 and 24 hpi (Figure 1F). This suggests that the NanoLuc insertion into the FCV-LC region impacts LC-associated viral egress.

To investigate the stability of FCV-Urbana^NL^ reporter virus, FCV-Urbana^NL^ was serially passaged in CRFK cells at a MOI 1 (Figure 2A). Virus was collected following each passage and titrated by plaque assay, before the passaged viruses were assessed for NanoLuc activity following infection of CRFK cells at 100 PFU/well for 16 hours. Viral titres increased significantly by passage 10, with the first significant increase observed between passages 3 and 4 (Figure 2B). This increase in infectious titre was mirrored by a decrease in NanoLuc activity at passage 4, followed by a significant decrease at passage 5 prior to reaching background levels of NanoLuc activity by passage 7, suggesting that the FCV-Urbana^NL^ no longer contained the NanoLuc sequence from LC by passage 4 (Figure 2C). To confirm the excision of the NanoLuc sequence, viral RNA was purified from each passage prior to the amplification of ORF2 by RT-PCR. Figure 2D shows the initial excision of the NanoLuc sequence beginning in passage 3 prior to removal of the NL sequence by passage 5, as highlighted by the reduction in size of the ORF2 amplicon from ∼2.3 kb to ∼1.8 kb.

**Figure 2:**
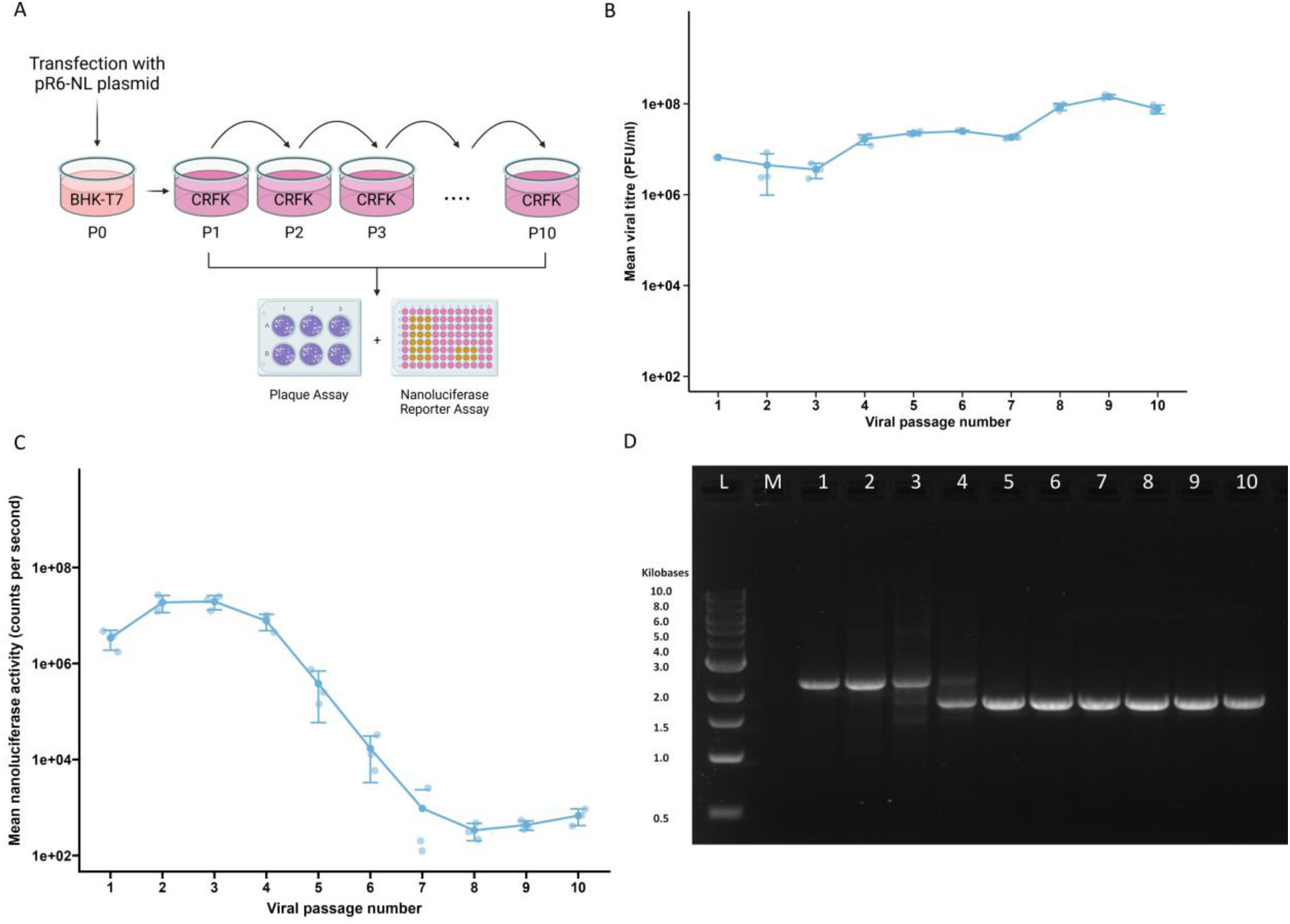
Stability of NL-FCV reporter virus. **(A)** Schematic of passaging workflow. FCV-NL plasmids were transfected into BHK-T7 cells to produce a P0 virus stock. The resultant virus was subsequently serially passaged 10 times in CRFK cells. **(B and C)** Viral titres were assessed by both NanoLuc assays measuring luminescence and plaque assays measuring viral titres (PFU/ml) at low MOI infections. (n=3) **(D)** Gel electrophoresis of RT-PCR analysis of NL insertion stability (L = Molecular weight ladder, M = Mock infected CRFK lysate, 1-10 = virus passage number).

### Chimaeric FCV-NanoLuc viruses infect a range of feline cell lines

Following the successful rescue of FCV-Urbana^NL^, we wanted to determine if it was possible to produce chimaeric capsid NanoLuc-FCV reporter viruses by replacing the FCV-Urbana^NL^ 3′ end ORF2 with the C-terminal end of LC and VP1 in ORF2 from other FCV viruses. To this end, we replaced the ORF2 region of FCV-Urbana^NL^ with the corresponding region of either FCV-F9, a commonly used vaccine strain, or FCV-NSW E1, a virulent systemic clinical isolate (10). These reporter viruses were termed FCV-F9^NL^ and FCV-NSW E1^NL^, respectively (Figure 3A). Both chimaeric reporter viruses were successfully rescued and, similar to FCV-Urbana^NL^, both showed reduced replication kinetics compared to the parent viruses, with FCV-NSW E1^NL^ displaying significant attenuation. Nevertheless, both viruses showed exponential growth kinetics (Figure 3B). We also evaluated the specificity and functionality of these reporter viruses using NanoLuc assays on CRFK cells. Similar to FCV-Urbana^NL^, both FCV-F9^NL^ and FCV-NSW E1^NL^ displayed significant increases in luminescence by 8 hpi, leading to a peak at 24 hpi, in contrast to the infections with the parental viruses that produced background levels of luminescence (Figure 3C).

**Figure 3:**
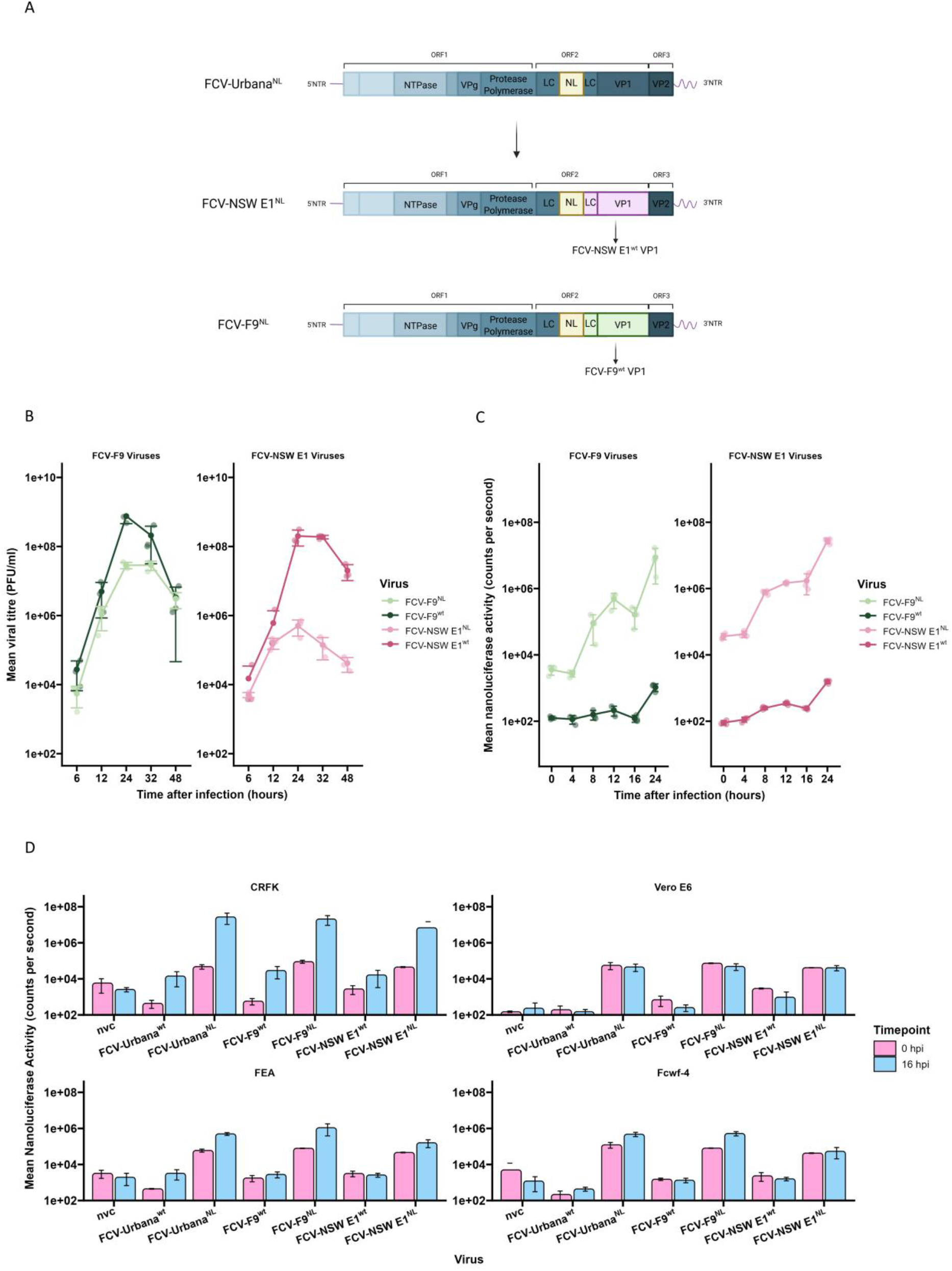
Infectivity of WT-FCV and NanoLuc-FCV strains in different cell lines. **(A)** Schematic depicting the generation of chimaeric FCV NanoLuc reporter viruses encoding the VP1 of FCV-F9 and FCV-NSW E1 **(B)** Comparison of viral replication kinetics between FCV-F9^wt^/^NL^ and FCV-NSW E1^wt/NL^ following infection at MOI 0.1. **(C)** NanoLuc activity elicited FCV-F9^wt^/^NL^ and FCV-NSW E1^wt/NL^ following infection at 100 PFU/well **(D)** FCV-Urbana^wt/NL^, FCV-F9^wt/NL^ and FCV-VS4^wt/NL^, were used to infect a range of feline-derived cell lines (CRFK, FEA and Fcwf-4) and Vero-E6 cells, at 100 PFU/well. Luminescence was measured at 0 and 16 hours after infection to evaluate the effectiveness of reporter viruses to infect various cell lines (n=3).

To evaluate the cell tropism specificity of our FCV^NL^ reporter viruses, their capacity to infect multiple feline-derived cell lines was assessed in parallel with their parental viruses. CRFKs, FEA and Fcwf-4 cells were infected at 100 PFU/well and luminescence was measured 0 and 16 hours after infection. Additionally, as a negative control, Vero-E6 cells were infected with 100 PFU/well, as Vero E6 cells are non-permissive for FCV (Figure 3D). NanoLuc activity increased for each FCV^NL^ virus in all feline-derived cells lines, whereas no observable difference in luminescence was detected following infection in Vero E6 cells. These data show that FCV^NL^ reporter viruses can effectively infect multiple feline cell lines and provide a sensitive readout, highlighting their potential for a range of applications in FCV research.

### Assessment of neutralising antibody activity using FCV^NL^

Traditional approaches to assess neutralising antibody activity often rely on plaque reduction neutralisation tests (PRNTs) (15). However, NanoLuc-Reporter viruses are equally effective, reducing assay time whilst providing an increased dynamic range (21–23). Therefore, to evaluate the potential of our FCV^NL^ reporter viruses to assess neutralising antibody activity, we tested three monoclonal antibodies (mAb) previously shown to target specific regions of FCV-Urbana VP1; mAb E5 targets a highly conserved non-neutralising epitope (PADGY) and mAb E10 targets an FCV-Urbana specific neutralising epitope (ITTANQY), whereas mAb C9 has been shown to target a neutralising conformational epitope on FCV-Urbana VP1 with unknown neutralising activity against other viral strains (24). We evaluated the neutralising activity for each mAb against our panel of FCV^NL^ reporter viruses (Figure 4). In line with previous findings, mAb E5 showed low levels of neutralising activity at high concentrations against FCV-Urbana^NL^, whilst mAb E10 strongly neutralised FCV-Urbana^NL^. However, mAb E10 did not neutralise FCV-F9^NL^ or FCV-NSW E1^NL^, confirming the specificity of mAb E10 for FCV-Urbana. Interestingly, mAb C9 exhibited strong neutralising activity against FCV-Urbana^NL^ but only low levels of neutralising activity at high concentrations against FCV-F9^NL^ and FCV-NSW E1^NL^, likely reflecting conformational differences in the epitopes recognised by mAb C9. These results demonstrate the utility of FCV^NL^ reporter viruses as tools to assess neutralising antibody activity.

**Figure 4:**
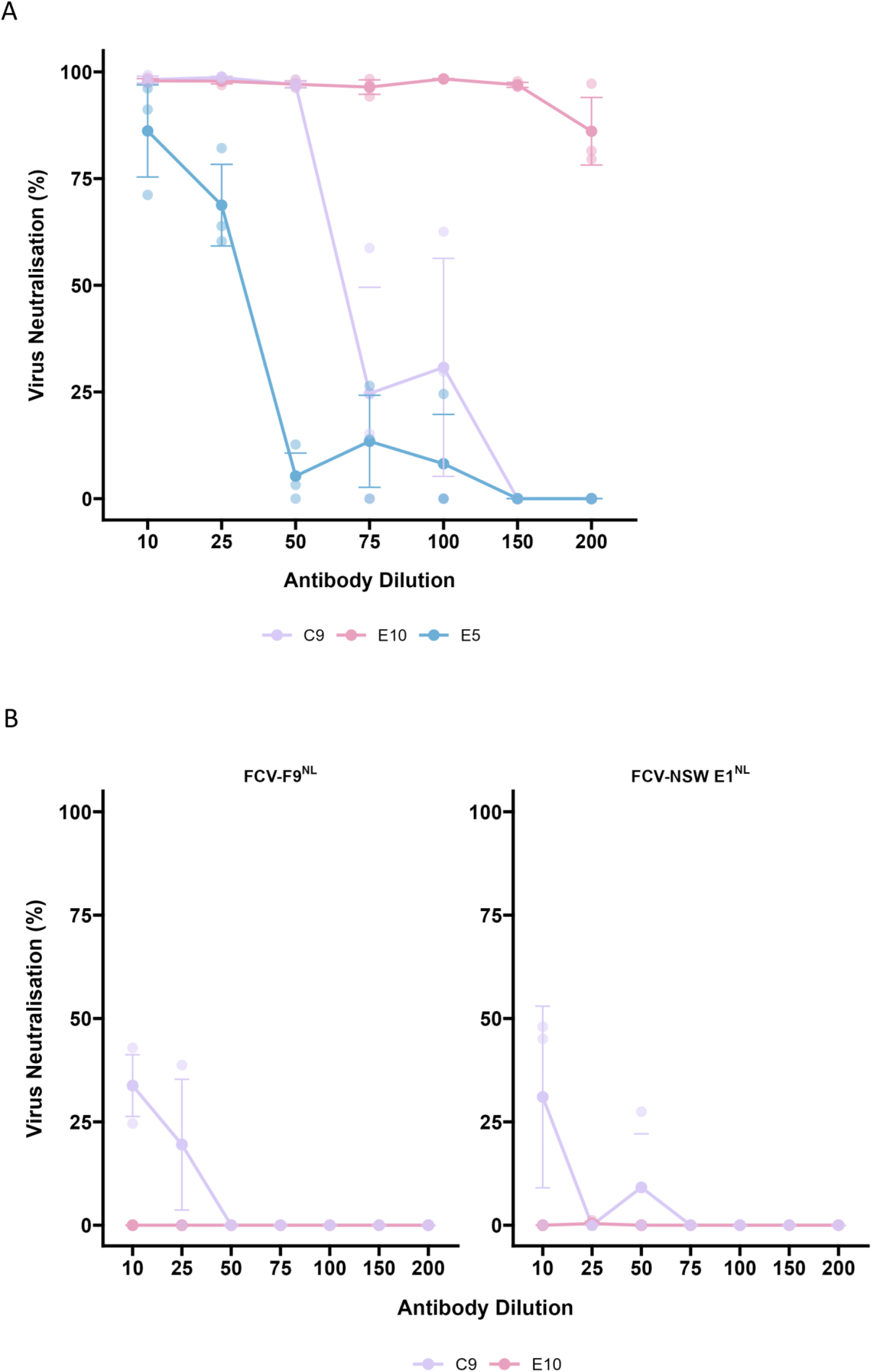
Utility of NanoLuc-FCV reporter viruses as a molecular tool to determine virus neutralisation. A panel of NanoLuc-FCV reporter viruses (FCV-Urbana^NL^, FCV-F9^NL^, FCV-NSW E1^NL^) were assessed in their activity against FCV-specific monoclonal antibodies, originally raised against FCV-Urbana^wt^. Neutralising (E10), conformational neutralising (C9) and non-neutralising (E5) antibodies were incubated with infectious virus, then plated onto cells at 100 PFU/well and incubated overnight; NanoLuc activity was measured 16-18 hours after infection. **(A)** Neutralising activity of mAbs against FCV-Urbana^NL^ **(B)** Neutralising activity of mAbs E10 and C9 against FCV-F9^NL^ and FCV-NSW E1^NL^ (n =3).

### Evaluation of antiviral compounds using FCV-NanoLuc

Reporter viruses also allow chemical inhibitors to be evaluated at a higher throughput than conventional viral titration approaches. Therefore, to test the utility of FCV^NL^ reporter viruses to assess antiviral compounds, we determined the inhibitory effect of four known antiviral compounds, namely ribavirin, nitazoxanide (NTZ) and 2′-C-methylcytidine (2′CMC), which have all been shown to inhibit FCV replication, and GS-441524 (Fig. 5). The latter is the active metabolite of the nucleoside analogue antiviral compound, remdesivir, now commonly used to treat cats with feline infectious peritonitis, caused by feline coronavirus (FCoV) infection (25,26). Cells were incubated with a range of concentrations of each antiviral for 1 hour prior to infection with 100 PFU/well FCV-Urbana^NL^ in the presence of antiviral and then incubated for 16 hours. Consistent with previous studies, all four compounds displayed antiviral activity associated with reductions in viral replication. NTZ exhibited the highest activity, with an IC_50_ of 1.77 μM, followed by 2′ CMC (IC_50_ = 6.59 μM). GS-441524 displayed moderate antiviral activity, producing an IC_50_ of 13.91 μM, whereas treatment with ribavirin led to a more modest reduction in replication, requiring higher concentrations to achieve antiviral activity (IC_50_ = 137.43 μM). Overall, these data demonstrate that FCV^NL^ reporter viruses can be used to identify and evaluate novel antiviral compounds.

**Figure 5:**
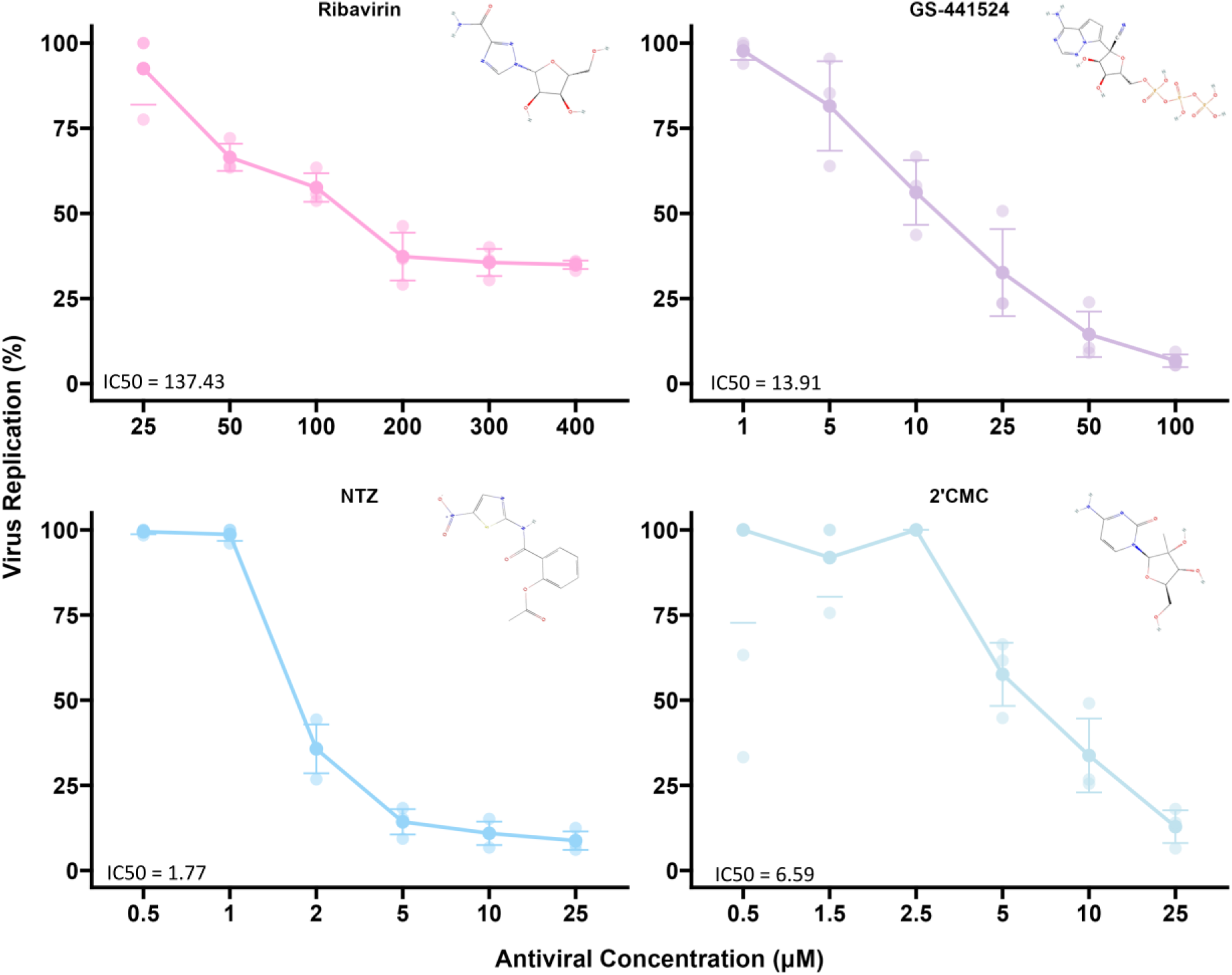
Utility of the NanoLuc-FCV reporter virus system in drug discovery. The antiviral activity of a panel of known antivirals was tested against FCV-Urbana^NL^: **(A)** Ribavirin, **(B)** NTZ, **(C)** 2′CMC and **(D)** GS-441524. CRFK cells were treated with each antiviral compound 1 hour prior to infection and the media supplemented with each compound for the duration of infection. At 16 hpi, luciferase activity was measured as a proxy for viral replication. The percentage of virus replication was normalised to the no inhibitor control in each assay (n=3). IC_50_ values are displayed as µM.

## Discussion

Despite the routine administration of FCV vaccines, FCV continues to pose a significant veterinary health burden (4). Therefore, there is a need for new molecular tools to improve our understanding of virus entry and replication and to facilitate the development of effective antiviral therapeutics, particularly considering the global emergence of VS-FCV strains. To address this need, we generated a genetically encoded, replication competent NanoLuc reporter virus for FCV. Reporter viruses are increasingly used in serological studies and antiviral drug screening, owing to their superior speed, sensitivity and scalability compared with conventional plaque reduction assays (15,27). Consistent with these advantages, FCV^NL^ viruses produced robust reporter signals within 8 hpi, and the assay dynamic range expanded by up to three orders of magnitude between 8-24 hpi (Figs. 1 & 3). By comparison, plaque reduction assays generally require 48 h for plaque formation and are limited by the relatively low number of plaques that can be accurately quantified per well. Together, these results demonstrated that FCV^NL^ viruses provide a rapid and highly sensitive platform for studying FCV replication and evaluating antiviral activity.

To generate our highly sensitive NanoLuc reporter virus, we built on the previous work of Abente *et al.* and replaced the fluorescent protein marker inserted into LC with NanoLuc. As previously reported for other FCV reporter viruses (20), we observed a reduction in the viral titre in comparison to the wild type virus (Figs. 1 & 3). Since it has been suggested that the LC protein of FCV plays a key role in viral egress through the induction of host cell permeabilisation (28–30), we tested whether the insertion of NanoLuc impacted viral release (Figs. 1E & 1F). Although freeze-thawing infected cells led to an increase in infectious virus, this did not account for differences in viral titre between the wild type and reporter viruses. These findings indicated that the addition of NanoLuc into the LC impacts both viral replication and viral egress.

The impact of introducing NanoLuc into LC mirrors observations from a recently reported murine norovirus (MNV) reporter virus (27), in which introduction of a HiBiT tag between NS4 and NS5 similarly attenuated viral replication kinetics. Reporter stability was comparable in both systems. FCV^NL^ displayed a significant reduction in NanoLuc activity by passage 4, followed by complete deletion of the NanoLuc gene by passage 5, whereas the MNV HiBiT virus retained only ∼10% NanoLuc activity by passage 4. Collectively, these data suggest that the utility of these reporter viruses is constrained by limited genetic stability upon serial passage. Consequently, reporter virus stocks should ideally only be used up to passage 3, to ensure robust and reproducible reporter virus activity. Notably, despite substantial differences in the size of the inserted sequences, the tolerance for these insertions appears to be broadly conserved across different caliciviruses.

Olson *et al.* suggested that the insertion of the HiBiT tag between NS4 and NS5, or other potential insertion sites, might be applicable to other caliciviruses. FCV belongs to the *Vesivirus* genus of caliciviruses, which is unique in being the only genus of calicivirus that expresses the LC protein, which can tolerate genetic insertions without completely attenuating the virus (20). An advantage of our genetically encoded reporter virus system is the ability to assess viral replication in any susceptible cell line using a simple lysis buffer as highlighted in Fig. 3. In contrast, the MNV HiBiT virus requires the addition of the complementary LgBiT as well as the lysis buffer to assess viral replication kinetics, increasing the cost of the assay.

Since a key application of reporter viruses is the quantification of the serological responses induced by viral infection or vaccination, we assessed the ability of FCV^NL^ to quantify the neutralising activity of previously characterised mAbs (24) (Fig. 4). Our data recapitulated the conclusions previously reported by Cubillos-Zapata *et al.*, with both mAbs E10 and C9 displaying over 90% neutralising activity at 1 in 150 and 1 in 50 dilutions, respectively. However, although mAb E5 displayed no neutralising activity at 1 in 20 dilution by PRNT, we observed over 50% neutralising activity at dilutions up to 1 in 25, highlighting the increased sensitivity of the reporter virus assay in comparison to conventional PRNT. We also investigated the breadth of mAb neutralisation by testing mAbs against reporter viruses containing heterologous VP1 proteins, assessing the capsid-specific neutralising activity in the same genetic background. Neither mAb E10 nor C9 displayed detectable neutralising activity against FCV-F9^NL^ nor FCV-NSW E1^NL^. The finding for mAb E10 was not unexpected, as this mAb targets a highly specific epitope in FCV-Urbana VP1, ITTANQY, at positions 445-451, with corresponding sequences in FCV-F9^NL^ and FCV-NSW E1^NL^ being ITTATGY and IATKDDY, respectively. However, mAb C9 targets an unknown conformational epitope on the FCV-Urbana capsid; therefore, we hypothesised that there could be cross-neutralisation between FCV-Urbana and both FCV-F9 and FCV-NSW E1 capsid proteins. Surprisingly, little cross-neutralising activity was observed, which might be explained by the high levels of conformational flexibility observed in calicivirus capsids, in addition to the genetic diversity displayed by the hypervariable region of FCV VP1 (31). Further studies are required to identify the conformational epitope targeted by mAb C9 to determine whether this antibody could neutralise isolates other than FCV-Urbana.

Reporter viruses allow the rapid identification and characterisation of novel antiviral compounds. Therefore, we assessed our FCV^NL^ reporter viruses against ribavirin, NTZ and 2′CMC, which all have antiviral activity against FCV, in addition to GS-441524, a compound now frequently used to treat cats with feline infectious peritonitis. Encouragingly, our results closely aligned with previously published data, with ribavirin displaying modest antiviral activity, and both NTZ and 2′CMC displaying lower micromolar IC_50_ values, at 1.77 and 6.59 μM respectively (10,32,33). Of particular interest, GS-441524 displayed pronounced antiviral activity at low micromolar concentrations, although it was less potent for FCV compared to FCoV, with IC_50_s of 13.91 μM and 0.78 μM, respectively (25). These data suggest that the FCV^NL^ reporter virus not only enables the identification and characterisation of novel antiviral compounds but also suggests that GS-441524 could represent an effective therapeutic option for cats infected with FCV as it does with FCoV. A key area for future study is to extend this methodology to produce NanoLuc reporter viruses in the context of complete VS-FCV genomes. This would provide the ability to rapidly assess phenotypical characteristics such as systemic virulence, tissue tropism and pathogenicity in addition to antiviral susceptibility in the context of clinically relevant isolates.

Overall, the data demonstrate that our FCV^NL^ reporter virus system is suitable for a range of applications, including viral replication studies, serological analyses and antiviral drug screening. Relative to conventional approaches such as PRNTs and immunofluorescent antibody tests, the reporter system offers several advantages, including reduced assay times and a substantially expanded dynamic range. Furthermore, the availability of NanoLuc substrates suitable for *in vivo* bioluminescence imaging raises the possibility of extending these applications to studies of viral pathogenicity. In conclusion, these findings establish FCV^NL^ as a versatile molecular tool with the potential to facilitate future investigations of FCV biology and improve our understanding of both FCV and VS-FCV infections.

## Methods

### Cell Culture and Viruses

Crandell Rees feline kidney (CRFK) cells were cultured in Dulbecco’s modified Eagles medium (DMEM, Thermo Fischer Scientific), which was supplemented with 10% (v/v) foetal bovine serum (FBS, Thermo Fischer Scientific), L-glutamine (2 mM) and penicillin (100U/mL)-streptomycin (100μg/mL) (complete DMEM). Feline embryonic kidney (FEA) cells, *Felis catus* whole foetus (Fcwf-4) cells (kindly provided by Susan Baker, Loyola University Chicago), and Vero-E6 cells were maintained as outlined above. Baby hamster kidney (BHK) cells expressing T7 RNA polymerase (BHK-T7) were cultured in DMEM supplemented with 5% FBS, L-glutamine (2 mM) and penicillin (100U/mL)-streptomycin (100μg/mL). Cells were passaged using 0.05% Trypsin-EDTA and maintained at 37℃ and 5% CO_2_.

FCV-Urbana was rescued using reverse genetics as described below. FCV-F9 was kindly provided by Wim Hesselink, Intervet. FCV-NSW E1 was kindly provided by Vanessa Barrs and Matteo Bordicchio (University of Sydney, Australia).

### Plasmid construction

A virus rescue plasmid (pR6) encoding the complete genome of wild-type FCV-Urbana was kindly provided by Dr Stanislav Sosnovtsev, NIH. This plasmid was modified to generate an FCV-Urbana plasmid containing a NanoLuc sequence. The *NLuc* gene was amplified by using the polymerase chain reaction (PCR) using a Forward primer to introduce a linker and the *KpnI* site (5′-GTTTAAACGGTACCGGCGGCAGCGTCTTCACACTC-3′) and a reverse primer to introduce a linker sequence at the 3′ of NLuc and the *AflII* restriction site (5′-CGAACGCATTCTGGCGTCCGGCGGCCTTAAGAATGAG-3′) from plasmid pNL1.1 (NanoLuc) (Promega), which was then inserted into the LC region of the pR6 FCV-Urbana^wt^ vector using *KpnI* and *AflII* (New England Biolabs (NEB)). The resulting pR6-NL plasmid was then transformed into DH5α (NEB 5-alpha) and purified by Maxi-prep kit (Qiagen) according to the manufacturer’s instructions.

To produce the chimaeric VP1-swap viruses, CRFK cells were infected with either FCV-F9 or FCV-NSW E1 at an MOI 1 for 6 hours. At 6 hpi, cells were lysed prior to total RNA purification using the Monarch® Total RNA Miniprep Kit (NEB). Following RNA extraction, cDNA was synthesised using SuperScript IV reverse transcriptase and oligo(dT) (Invitrogen) as per the manufacturer’s instructions. The ORF2 region was amplified from nucleotide position 5578 into the first 4 amino acids of ORF3 at nucleotide position 7328 and inserted into pR6-NL using *AflII* and *AvrII* (NEB). The resulting pR6-NL F9 and pR6-NL NSW-E1 plasmids were transformed and amplified as described above. All PCR steps were carried out with Phusion polymerase (NEB) using the manufacturer’s guidelines with verification via Sanger sequencing (Eurofins).

### Transfection and virus rescue

To rescue virus, BHK-T7 cells were seeded at ∼80% confluency in 12-well plates. Individual wells were transfected with 2 μg of virus rescue plasmid (pR6, pR6-NL, pR6-NL F9 or pR6- NL NSW E1) alongside 0.5 μg pCAGGS-T7 expression plasmid to boost the initial round of viral transcription. Briefly, plasmids were mixed with 250 μL of pre-warmed Opti-MEM and 7.5 μL Mirus TransIT-LT1 transfection reagent (Cambridge Biosciences) and incubated at 25℃ for 18 minutes as per the manufacturer’s instructions. The BHK-T7 cells were washed with serum-free antibiotic-free DMEM, supplemented with 2mM L-glutamine prior to incubation in 700 μL serum-free antibiotic-free DMEM. Transfection mixtures were added to cells drop-wise prior to incubation at 37°C/ 5% CO_2_ overnight.

24 hours post-transfection, the transfected BHK-T7 cells were overlaid with 4 x 10^5^ CRFK cells resuspended in serum-free DMEM (supplemented with L-glutamine (2 mM) and penicillin (100U/mL)-streptomycin (100μg/mL)). The transfected cells were then cultured for a further 48 hours, at 37°C/ 5% CO_2_. At 72 hours post-transfection, BHK-T7-CRFK co-cultures were freeze-thawed three times to enhance virus particle release. Cell lysates were centrifuged at 2400 *g* for 5 minutes to remove cell debris. Following centrifugation, the supernatant, determined as Passage 0 (P0), was collected and frozen at-80°C.

To obtain P1 stocks of FCV-Urbana^wt^ and the FCV^NL^ viruses, confluent T25 flasks containing CRFK cells were washed with PBS and incubated with the P0 lysates. Following infection, the CRFK cells were incubated for 2 hours at 37°C/ 5% CO_2_. The infection media was then aspirated and replaced with fresh sf-DMEM. Cells were then incubated further at 37°C/ 5% CO_2_, until a 90-100% cytopathic effect (CPE) was observed. Viral supernatants were then harvested, freeze-thawed three times to enhance the release of virus particles, aliquoted and stored at-80℃. If CPE failed to appear by day 5 after inoculation, the virus rescue was considered unsuccessful.

To produce P2 virus stocks, where required, confluent CRFK cells were infected at a MOI of 0.1 or 1, for wild-type viruses or NanoLuc containing viruses, respectively. Once the infection reached 90-100% CPE, cells were freeze-thawed three times to aid the release of viral particles, and viral supernatant were aliquoted and stored at-80℃ until titration.

### Virus Titration

Viral titres were determined by plaque assay. Briefly, confluent 6-well plates containing CRFK cells were washed with sf-DMEM prior to infection. Virus stocks were serially diluted in 10- fold dilutions (10^-1^-10^-8^) in sf-DMEM and 800 µL of dilutions 10^-3^-10^-8^ were added to cell monolayers and incubated at 37°C/ 5% CO_2_ for 2 hours. Following incubation, infected cell monolayers were overlaid with 2 mL 1.2% Avicel (w/v) diluted in sf-DMEM, leading to a final concentration ∼0.9% Avicel (w/v). Plaque assays were then incubated for 32 hours at 37°C/ 5% CO_2_.

After 32 hours, cells were fixed by adding 2 mL 10% formaldehyde (v/v) to each well and incubated at room temperature for 30 minutes. The formaldehyde-Avicel overlay was then aspirated from wells and then cells were washed with PBS and stained with Coomassie Blue. Following staining, plaques were counted, and the viral titre was calculated in plaque forming units/mL (PFU/mL).

### FCV-NanoLuc sequence validation

To validate FCV-NanoLuc viruses and investigate their stability, FCV-Urbana^NL^ was serially passaged 10 times in CRFK cells at a MOI 1 with viral supernatants collected and stored after each passage. To obtain viral RNA, confluent CRFK cells seeded in 6-well plates were infected at a MOI of 1 in sf-DMEM and incubated at 37°C/ 5% CO_2_ for 6 hours. At 6 hpi, cells were lysed prior to total RNA purification using the Monarch® Total RNA Miniprep Kit (NEB). Following RNA extraction, cDNA was synthesised using SuperScript IV reverse transcriptase and oligo(dT) (Invitrogen) as per the manufacturer’s instructions. To determine the stability of the NanoLuc reporter gene, we designed a forward primer which binds upstream of the NanoLuc insert in LC (5′-GCGATGATGAGTGGTCTTC-3′) and a reverse primer (5′- GTAACAGTATCAATCAAGCCTAGG-3′) which amplifies from the 3′ end of ORF2. Following PCR, we determined the presence of the NanoLuc reporter gene through 1% agarose gel electrophoresis.

### NanoLuc assay

To determine NanoLuc activity of our reporter virus as a proxy for viral replication, confluent 96-well plates seeded with CRFK cells were infected with 100 PFU/well in sf-DMEM. Briefly, cells were aspirated and overlaid with the viral inoculum, the cells were incubated at 37°C/ 5% CO_2_ for 16-18 hours. Subsequently, a NanoLuc lysis buffer and substrate solution (Promega) was prepared to the manufacturer’s instructions and added directly to each well. Plates were then read using an EnSight Multimode plate reader (Revvity) to measure NanoLuc activity. NanoLuc reporter assays using other FEA, Fcwf-4 and Vero E6 cells were performed as outlined above.

### Viral neutralisation and antiviral assays

To assess the neutralising activity of monoclonal antibodies (mAbs) raised against FCV Urbana, we used the NanoLuc assay described above with additional amendments described below.

Virus stocks were diluted to 200 PFU/well, while the Abs were diluted to twice the desired final concentration. The mAbs were then diluted 1 in 2 in 12 well plates when added to the diluted virus stocks and incubated at 37°C/ 5% CO_2_ for one hour to allow virus-antibody binding to occur. Following incubation, sf-DMEM was aspirated from the CRFK cells and the virus-mAb mixture was used to inoculate cells. Plates were then incubated for 16-18 hours at 37°C/ 5% CO_2_. A NanoLuc lysis buffer and substrate solution (Promega) was then prepared and added to infected wells as per the manufacturer’s instructions. Plates were then read using an EnSight Multimode plate reader (Revvity) and viral replication activity was calculated.

To determine the antiviral activity of a range of compounds, we used the NanoLuc assay described above with the additional amendments described here.

Antiviral compounds were diluted in sf-DMEM to a range of concentrations as follows:

-GS-441524 (Fisher Scientific): 100 μM, 50 μM, 25 μM, 10 μM, 5 μM and 1 μM
- Ribavirin (Merck): 400 μM, 300 μM, 200 μM, 100 μM, 50 μM and 25 μM
- Nitazoxanide (NTZ) (Merck): 25 μM, 10 μM, 5 μM, 2 μM, 1 μM and 0.5 μM
- 2′-C-Methylcytidine (2’CMC) (Merck): 25 μM, 10 μM, 5 μM, 2.5 μM, 1.5 μM and 0.5 μM

Antiviral compounds were then incubated plated with cells in triplicate at the concentrations above and incubated for an hour at 37°C/ 5% CO_2_. After incubation, antivirals were aspirated from cells and virus stocks diluted to 100 PFU/μL were used to inoculate cells. Plates were incubated for 16-18 hours at 37°C/ 5% CO_2_. A NanoLuc lysis buffer and substrate solution (Promega) was then prepared and added to infected wells. Plates were then read using an EnSight Multimode plate reader and viral titres were calculated.

## Statistical Analysis

Data panels were prepared and analysed with RStudio (version 4.5.1) and Microsoft Excel (version 2408) with statistical methods described below. Statistical significance was set as P < 0.05. Figure 1F data are shown as mean ± SD (standard deviation) of triplicates. An unpaired Student’s t-test was performed to compare viral titres of FCV-Urbana^wt^ and FCV- Urbana^NL^ following freeze-thaw at each time-point, where * denotes P < 0.05 and ns means no significant differences.

## Authors and Contributions

Conceptualisation: LS, MJH, BJW. Funding acquisition: MJH, BJW, DB, LS. Investigation: LS, HS, KU, RA, MM, LHT, SO. Resources: JB, SB, VB, MB Supervision: MJH, BJW, WW and LS. Writing – original draft: HS, LS and MJH. Writing – review and editing: LS, HS, JB, WW, BJW and MJH.

## Acknowledgments

We thank other members of the Hosie/Willett group, at the MRC-University of Glasgow Centre for Virus Research, for their insightful contributions.

